# Macromolecular tissue volume mapping of lateral geniculate nucleus subdivisions in living human brains

**DOI:** 10.1101/2020.12.26.424373

**Authors:** Hiroki Oishi, Hiromasa Takemura, Kaoru Amano

## Abstract

The human lateral geniculate nucleus (LGN) is composed mainly of the magnocellular and parvocellular subdivisions. The non-invasive identification of these subdivisions is, however, difficult due to the small size of the LGN. Here we propose a method to identify these subdivisions by combining two structural MR measures: high-resolution proton-density weighted images and macromolecular tissue volume maps. We collected MRI data from 15 healthy subjects and found that the spatial organization of these subdivisions identified by our method was consistent with post-mortem histological data. Furthermore, the stimulus selectivity in these regions measured by functional MRI was consistent with physiological studies. These results suggest that macromolecular tissue volume mapping is a promising approach to evaluating the tissue properties of LGN subdivisions in living humans. This method potentially will enable neuroscientific and clinical hypotheses about the human LGN to be tested.

## Introduction

The lateral geniculate nucleus (LGN) plays an essential role in human visual processing by receiving visual inputs from retinal ganglion cells and transferring those signals to the primary visual cortex (Nassi and Callaway, 2009). Besides its function as a relay station, a growing body of evidence suggests that the LGN is involved in a wide range of visual functions, including eye-specific dominance and suppression during binocular rivalry (Haynes et al., 2005), visual attention (Ling et al., 2015; O’Connor et al., 2002; Schneider and Kastner, 2009), and visual perceptual learning (Yu et al., 2016). The LGN is also involved in the neuronal synchrony widely observed in the visual cortex (Hughes et al., 2004; Liu et al., 2012; Minami et al., 2020).

Human LGN is composed of six layers that are categorized into two major subdivisions, the magnocellular (M) and parvocellular (P) subdivisions. These subdivisions are clearly distinguishable on the basis of cell size (Hickey and Guillery, 1979; Gupta et al., 2006). A number of studies have suggested that the M and P subdivisions have complementary roles in visual processing by demonstrating distinct spatial, temporal, luminance, and chromatic stimuli preferences (Denison et al., 2014; Derrington and Lennie, 1984; Schiller et al., 1990; Usrey and Reid, 2000). According to psychophysical performances in response to specific types of visual stimuli and neural responses in the M and P subdivisions, these two subdivisions have been proposed to have distinct roles in attention (Yeshurun and Levy, 2003) and reading (Demb et al., 1998; Main et al., 2014; Stein, 2001). A psychophysical study using a motion coherence task also led to a hypothesis that the M subdivision is damaged earlier than the P subdivision as a consequence of glaucoma (Joffe et al., 1997). However, the hypotheses proposed by previous psychophysical studies remain speculative, because they are derived mostly from similarities between the neural responses of the M and P subdivisions and stimuli-dependent psychophysical performances without assessing the differences in functional and structural measurements between the two. Therefore, it is essential to establish non-invasive measurements to enable the direct comparison of neuroimaging data from the LGN subdivisions and psychophysical data, while also comparing the properties of the M and P subdivisions with those of other visual cortical areas.

Non-invasive neuroimaging-based measurements of the structural properties of human LGN subdivisions have not been established, because of the requirement for high-resolution, quantitative MRI measurements for the LGN, which has a small volume around 180 mm^3^ (Mcketton et al., 2014). While functional MRI (fMRI) has been used to localize M and P subdivisions based upon the blood oxygenation level-dependent (BOLD) response selectivity for distinct visual stimuli (Denison et al., 2014; Zhang et al., 2016, 2015), the spatial resolution (1.25 × 1.25 × 1.2 mm^3^–1.75 × 1.75 × 1.5 mm^3^) and robustness of the measurements have been limited. fMRI measurements also require the use of visual stimuli, which limit the applicability for clinical populations with visual field loss.

To examine the properties of LGN subdivisions in individual living humans, here we propose a method by which the subdivisions can be identified using structural MRI measurements. To this end, we primarily used macromolecular tissue volume (MTV) mapping, which is a quantitative structural MRI method that is sensitive to a fraction of non-water macromolecules (Mezer et al., 2013). We hypothesized that the MTV fraction would enable the M and P subdivisions to be distinguished as (1) MTV is sensitive to lipids which constitute cell membranes and myelin (Filo et al., 2019; Mezer et al., 2013; Shtangel and Mezer, 2020) and (2) neuronal cell and myelin densities differ between the M and P subdivisions (Hassler, 1966; Pistorio et al., 2006; Yücel et al., 2003, 2000). The first step in our proposed method was to identify the location and contour of the whole LGN using high-resolution proton-density (PD)-weighted imaging as employed in previous studies on human LGN (Giraldo-Chica and Schneider, 2018; Mcketton et al., 2014; Viviano and Schneider, 2015). We then defined the LGN subdivisions using MTV fraction data and the anatomically known volume ratio of the M and P subdivisions. Finally, we tested the validity of the definition based on (1) comparisons with histology data, (2) fMRI measurements of stimulus selectivity, and (3) an analysis of the test-retest reliability.

Using our newly developed method, we found a gradual change in the MTV fraction within the LGN along each axis (lateral-medial, dorsal-ventral, and anterior-posterior). This pattern of change was consistent among subjects, enabling the parcellation of the LGN into two subdivisions in a consistent manner with post-mortem human data. Moreover, the difference in stimulus selectivity of the BOLD response between the subdivisions identified by MTV was consistent with previous physiological studies. The MTV-based LGN parcellation was robust over measurements taken on different days. The parcellation using widely used non-quantitative methods such as the T1 weighted/T2 weighted (T1w/T2w) ratio map was not sufficient to identify the LGN subdivisions compared with MTV, suggesting that quantitative structural mapping is crucial to identifying the M and P subdivisions in human LGN. This study provides a novel method of non-invasively investigating the properties of LGN subdivisions in living human brains, which can be combined with functional or behavioral experiments to test neuroscientific or clinical hypotheses.

## Materials and Methods

### Subjects

Fifteen healthy volunteers (7 females; mean age, 23.53 years; standard deviation, 1.71 years; range, 21–26 years) participated in the study. All of the subjects had normal or corrected-to-normal vision with no clinical history of eye disease. All subjects provided written informed consent to take part in this study. The study was conducted in accordance with the ethical standards of the Declaration of Helsinki and approved by the local ethics and safety committees at the Center for Information and Neural Networks (CiNet), National Institute of Information and Communications Technology (NICT).

### Structural MRI data acquisition

All MRI data were collected at the CiNet using a 3T MAGNETOM SIEMENS Prisma scanner (Siemens Healthcare, Erlangen, Germany) with a 32-channel head coil.

#### T1 weighted MRI data acquisition

T1 weighted magnetization prepared-rapid gradient echo (MP-RAGE) images (voxel size, 0.75 mm × 1.0 mm × 0.75 mm; repetition time [TR], 1900 ms; echo time [TE], 3.58 ms; flip angle, 9°; matrix, 256 × 256; in-plane acceleration factor, 2) were acquired from all subjects. These images were used as references on which to co-register the subsequent MRI data (PD weighted, MTV, and fMRI data) in the same coordinate space for each individual subject. The total time for the T1 weighted MRI data was approximately 15 min for each subject.

#### Proton density (PD)- weighted MRI data acquisition

PD weighted images were acquired from all subjects to locate the LGN. Acquisition parameter of PD weighted images followed those used in a previous study (Viviano and Schneider, 2015; voxel size, 0.75 mm × 0.75 mm × 1.0 mm; TR, 3000 ms; TE, 21.0 ms; flip angle, 120°; matrix, 256 × 256; in-plane acceleration factor, 2). These images were acquired at least 40 times in all subjects. To improve the signal-to-noise ratio, we then continued to repeat the PD-weighted image acquisition if the subjects agreed (maximum number of repetitions: 60). Each image consisted of 50–60 coronal slices (slice thickness, 1 mm; no gap) covering the whole posterior thalamus. The total acquisition time for the PD weighted MRI data was approximately 60–90 min for each subject, depending on the number of repetitions.

#### MTV data acquisition

The MTV data were acquired from all subjects according to a previously described protocol (Mezer et al., 2013; Minami et al., 2020; Oishi et al., 2018; Takemura et al., 2019). Four fast low-angle shot (FLASH) images were measured with flip angles of 4°, 10°, 20°, and 30° (TR, 12 ms; TE, 2.43 ms) with 1 mm isotropic voxels. For the purposes of removing field inhomogeneities, five additional spin echo inversion recovery (SEIR) scans were also measured with an EPI readout (TR, 3 s; TE, 49 ms; 2 × acceleration). The inversion times were 50, 200, 400, 1200, and 2400 ms. The in-plane resolution and slice thickness of the additional scan were 2 × 2 mm^2^ and 4 mm, respectively. The total acquisition time for MTV data was approximately 35 min for each subject. For 11 subjects, we acquired MTV data again on a different day to evaluate the test-retest reproducibility.

#### T1w/T2w MRI data acquisition

We also acquired data from 13 subjects for a T1w/T2w ratio map, a technique widely used in the analysis of Human Connectome Project data (Glasser and Van Essen 2011). The T1 weighted image was acquired using a 3D MP-RAGE (TR, 2400 ms; TE, 2.06 ms; TI, 1000 ms; flip angle, 8°; bandwidth, 220 Hz/pixel; echo spacing, 7.5 ms; field of view, 256 mm × 256 mm × 176 mm; matrix, 256 × 256 × 176; voxel size, 1 mm isotropic resolution) sequence. The T2 weighted image was acquired using sampling perfection with application optimized contrast using different angle evolutions (SPACE: TR, 3200 ms; TE, 438 ms; flip angle, 120°; bandwidth, 574 Hz/pixel; echo spacing, 3.88 ms; turbo factor, 139; field of view, 256 mm × 256 mm × 176 mm; matrix, 256 × 256 × 176; voxel size, 1 mm isotropic resolution) sequence. While these acquisition protocols aimed to follow those used in the Human Connectome Project (Glasser and Van Essen 2011), they were not identical due to hardware differences. These data were collected using prescan normalization to reduce image intensity bias.

### Structural MRI data analysis

#### T1 weighted MRI data

The T1 weighted MRI images of individual subjects were interpolated and aligned to the ICBM 152 2009b symmetric template in the MNI152 database (http://www.bic.mni.mcgill.ca/ServicesAtlases/ICBM152NLin2009; isotropic resolution, 0.5 mm; Fonov et al., 2011, 2009) using a rigid-body transformation implemented in the FSL FLIRT tool. No spatial smoothing or normalization was performed. The T1 weighted MRI image aligned onto the MNI152 space was used for subsequent comparisons between MTV, fMRI, and histological data from individual brains in MNI coordinates.

#### PD weighted MRI data

The PD weighted image from the first scanning session was used as a reference, and all subsequent PD images were co-registered to the reference using a rigid-body transformation implemented in FSL. We then averaged all of the PD weighted images aligned to the first PD weighted image. The resulting averaged PD weighted image was then interpolated and aligned to the T1 weighted MRI data, which was aligned with the MNI coordinates at 0.5 mm isotropic resolution for subsequent analyses.

#### Quantitative MRI data

Using the mrQ software package (https://github.com/mezera/mrQ) in MATLAB, the FLASH and SEIR scans were processed to produce the MTV maps (Mezer et al., 2016, 2013). MTV quantifies the macromolecular tissue volume density by estimating a quantitative PD map from the FLASH images after correcting for RF coil bias by the mrQ analysis pipeline using SEIR-EPI scans (Barral et al., 2010; Mezer et al., 2013). Since cerebrospinal fluid (CSF) voxels are entirely filled with water, we assumed that they had a full water volume fraction (WVF). We then calculated the WVF ratio in cortical gray or white matter voxels in comparison with the CSF. The MTV was defined as follows: MTV = 1 − WVF, and this was used to quantify the non-proton macromolecule volume fraction in each voxel. Finally, the MTV map was aligned to the T1 weighted MRI data to enable further comparisons with other images in the same coordinate space. The full analysis pipeline can be found in previous publications (Mezer et al., 2016, 2013; Minami et al., 2020; Oishi et al., 2018; Takemura et al., 2019).

#### T1w/T2w MRI data

We obtained a T1w/T2w ratio map by co-registering a T2-weighted image to T1-weighted image using the FLIRT tool in FSL (Jenkinson et al., 2002) with six parameters (rigid body) and calculated the ratio between them. The T1w/T2w ratio map was then co-registered to the reference T1 weighted MRI data at the MNI coordinates.

### Identifying a LGN region-of-interest from PD-weighted MRI data

With reference to previous studies (Viviano and Schneider, 2015), we identified the position of the LGN in individual subjects based on the PD weighted image averaged across multiple acquisitions (Figure 1). We manually delineated the whole LGN based on visible intensity differences between the LGN and neighboring tissues (surrounding white matter and CSF) using the ITK-snap tool (http://www.itksnap.org/; Figure 1B). Delineation was performed in a series of coronal sections of PD weighted image because the coronal sections have the highest spatial resolution in comparison with the axial and sagittal sections. The whole LGN region-of-interest (ROI) was used in subsequent analyses to classify the M and P subdivisions using MTV.

**Figure 1.**
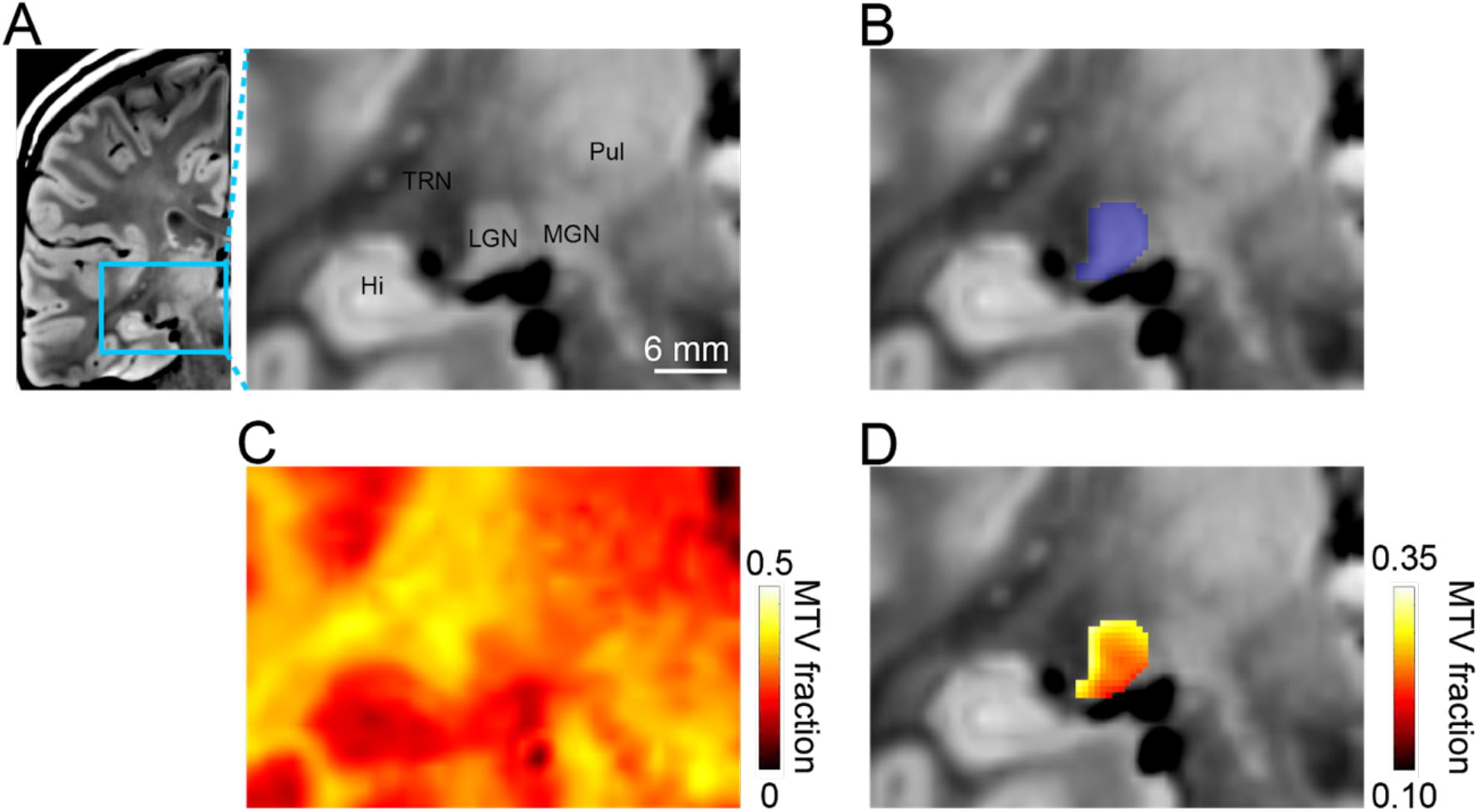
Procedure for identifying the whole LGN and LGN subdivisions in a single human subject from structural MRI data. (A) A coronal section of a proton density (PD) weighted image in a representative subject (left hemisphere, Subject S10). *Left panel,* the coronal PD weighted image of the whole left hemisphere. The cyan rectangle indicates the region magnified in the right panel. *Right panel,* the magnified PD weighted image near the LGN. TRN, thalamic reticular nucleus; Hi, hippocampus; MGN, medial geniculate nucleus; Pul, pulvinar. The scale bar (white line) indicates 6 mm. (B) The ROI covering the whole LGN (translucent blue), which was manually defined from the PD weighted image. (C) Macromolecular tissue volume (MTV) map co-registered with the PD weighted image. The hot color map corresponds to MTV fractions in individual voxels. (D) MTV fractions within the LGN ROI. The MTV fraction gradually changed along the superior-inferior and lateral-medial axes. Note that the scale of the MTV fraction differs from that of panel C.

### Parcellation of the LGN based on MTV and other structural MRI maps

We first rank-ordered all of the voxels within the whole LGN ROI based on their MTV fractions. Previous histological studies have identified that the P subdivision has a higher neuronal cell density (Hassler, 1966; Yücel et al., 2003, 2000) and greater myelin content (Pistorio et al., 2006) than the M subdivision, and therefore we hypothesized that the P subdivision would demonstrate larger MTV fractions than the M subdivision. Hence, we classified the 20% of voxels with the lowest MTV fraction as the putative M subdivision and the remaining 80% of voxels as the putative P subdivision (Figure 2B). This ratio was based on previously reported volumes of the LGN subdivisions from prior human histological studies (Andrews et al., 1997; Selemon and Begovic, 2007) and was used in a previous fMRI study (Denison et al., 2014). Figure 2 provides examples of MTV-based LGN parcellation in representative hemispheres.

**Figure 2.**
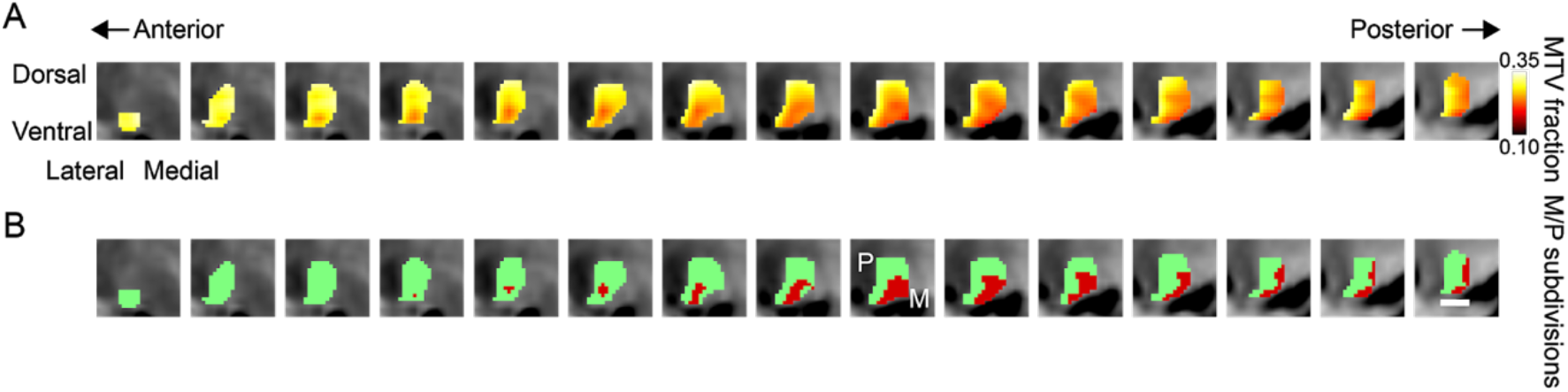
LGN subdivisions parcellated by MTV fraction on a series of coronal sections in a single representative hemisphere. (A) MTV fractions in the LGN ROI overlaid on a series of coronal sections of a PD weighted image (the left and right panels represent anterior and posterior sections; distance between sections: 0.5 mm) in a representative hemisphere (left hemisphere, subject S10). The conventions were identical to those used in Figure 1D. (B) The M and P subdivisions estimated from the MTV fractions in the hemisphere shown in panel A. We classified the 20% of voxels with the lowest MTV fractions as belonging to the M subdivision (dark red) and the remaining 80% of voxels as belonging to the P subdivision (light green). The scale bar indicates 4 mm.

Next, we attempted to parcellate the LGN based on the image intensity of the non-quantitative structural MRI maps (PD weighted image and T1w/T2w ratio map). For the PD-weighted image, we classified 20% of voxels with the highest image intensity as the putative M subdivision and the remaining 80% of voxels as the putative P subdivision, since the image contrast of PD weighted images demonstrates an opposite trend to that of MTV maps. For the T1w/T2w ratio map, we used the identical criteria as employed for the MTV mapping.

### Histological data (BigBrain) analysis

To compare the MTV-based parcellation of the LGN subdivisions with the histological definition, we analyzed the publicly available BigBrain data (100 μm version of the BigBrain 3-D Volume Data Release 2015 in MNI space from https://bigbrain.loris.ca; Amunts et al., 2013). In brief, BigBrain is a 3D reconstruction of 7404 histological sections of one post-mortem human brain that provides high-resolution anatomical data aligned with MNI coordinate space. In this database, all of the six layers in the human LGN are visible (see Figure 2-S3).

Manual segmentation of the M (layers 1–2) and P subdivisions (layers 3–6) of the human LGN was carried out on the BigBrain data. The manual segmentation was performed by a rater who was not an author of this work and unaware of the purpose of this study. We used this M and P subdivision definition from BigBrain as a reference with which to compare the MRI-based parcellation. An example of manual segmentation of the BigBrain data is shown in Figure 2-S3.

### Comparison between MRI data with histological data

To quantify the spatial organization of the M and P subdivisions, we calculated the spatial centers of both subdivisions in MRI and histological data, following the analysis used in a previous fMRI study (Denison et al., 2014). The 3D spatial centers of the M and P subdivisions were defined as the mean voxel coordinates in each spatial dimension (anterior-posterior, dorsal-ventral, and left-right) in MNI space. When co-registering the reference (T1 weighted) image to MNI space, the human thalamic nuclei showed individual differences in their positions, volume, and shape (Csernansky et al., 2004). Therefore, the position of the LGN and its subdivision in the MNI coordinates is variable among individual brains. Thus, we calculated the relative position of the center of each LGN subdivision with respect to the widths of the LGN in each spatial dimension to compare the LGN subdivisions between datasets and brains (Denison et al., 2014).

### Functional MRI data acquisition

All subjects took part in an additional fMRI experiment to investigate the stimulus selectivity of BOLD response in LGN voxels. We acquired fMRI data with 1.5 mm isotropic voxels for 10 subjects (S1–S10) and 2 mm isotropic voxels for 5 subjects (S11–S15).

#### Acquisition parameters

The fMRI data were acquired with an interleaved T2* weighted gradient echo sequence at an isotropic voxel size of 1.5 or 2.0 mm isotropic by using simultaneous multi-slice EPI sequences (TR, 2250 ms; TE, 40 ms; flip angle, 75°; in-plane field of view, 192 × 192 mm) provided by the Center for Magnetic Resonance Research, Department of Radiology, University of Minnesota (https://www.cmrr.umn.edu/multiband/; Moeller et al., 2010). Transverse axial slices (57 slices for 1.5 mm isotropic voxels and 50 slices for 2.0 mm isotropic voxels) with no gaps were oriented to cover the LGN and occipital lobe. Certain parameters differed for the acquisition of 1.5 mm and 2 mm isotropic voxels acquisitions (multi-band factor, 3; acquisition matrix, 128 × 128; echo spacing, 0.93 ms; partial Fourier, 6/8 for 1.5 mm isotropic voxels; multi-band factor, 2; acquisition matrix, 96 × 96; echo spacing, 0.68 ms; partial Fourier was not applied for 2 mm isotropic voxels), while other parameters were identical.

#### Stimuli, block design, and task

All visual stimuli were generated using Psychtoolbox 3 in MATLAB (http://psychtoolbox.org/). Stimuli were projected from a projector (WUX5000, Cannon) located outside the scanner room and reflected via a mirror onto a gamma-corrected translucent screen positioned over the subject’s head. Gamma-correction was applied using Mcalibrator2 (Ban and Yamamoto, 2013; https://github.com/hiroshiban/Mcalibrator2). Stimuli were presented on a full flat screen (416 mm × 222 mm) at a spatial resolution of 1920 × 1200 and a frame rate of 60 Hz. The screen was viewed via a mirror mounted over the subject’s eyes. The viewing distance and visual angle of the screen was 92 cm and 41.2° × 25.8°, respectively.

We adapted two types of stimuli (M-type and P-type stimuli; Figure 4A) designed to elicit selective BOLD responses in the M and P subdivisions (Denison et al., 2014; https://github.com/racheldenison/MPLocalizer). The M-type stimulus was a 100% contrast, black-white grating with low spatial frequency (0.5 cycles per degree) and higher flicker frequency (15 Hz). The P-type stimulus was a near-isoluminant red-green grating with higher spatial frequency (2 cycles per degree) and lower flicker frequency (5 Hz). The orientation of the grating (0°, 30°, 60°, 90°, 120°, or 150°) was changed every 3 s in a random manner. Prior to the fMRI experiment, we adjusted the luminance of the P-type stimuli to make it perceptually isoluminant using a flicker method (Ives, 1912; Minami and Amano, 2017).

We used a block design for the fMRI experiment, in which each run comprised 15 blocks (six blocks for each of the M- and P-type stimuli and three blocks with a blank screen). Each block was 20.25 s in duration (including 18 s for stimulus presentation and 2.25 s for the blank period, during which the subjects provided their responses). During each block, subjects were instructed to count the number of randomly presented targets, two-dimensional Gaussian contrast decrements within the stimuli, while maintaining fixation. During the blank period, the subjects reported how many targets they had seen during the previous stimuli block by pressing a button. The subjects completed 7–8 runs. This procedure and other details of the stimuli, task, and block design have been described previously (Denison et al., 2014).

### Functional MRI data analysis

The fMRI data were analyzed using mrVista (https://github.com/vistalab/vistasoft). We registered the data onto T1 weighted MRI data to enable comparisons with other MRI datasets. We corrected the slice timing to match the multi-slice acquisition order. The data were then corrected for the subject’s motion within and between scans. We fitted a general linear model consisting of predictors (M- and P-type stimuli were regressors) convolved with the hemodynamic response function (HRF; Boynton et al., 1996) to the time course of each voxel. We used the Boynton HRF to match the procedure in a previous fMRI study (Denison et al., 2014). By fitting the HRF model to the time series of BOLD responses, we estimated the beta values for the M- and P-type stimuli. We then calculated the difference between them (beta_M-P_) as follows:

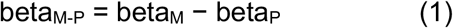

We used beta_M-P_ as an index for evaluating the stimulus selectivity of the LGN subdivisions identified by MTV. We averaged the beta_M-P_ across all voxels in each LGN subdivision parcellated on the basis of MTV maps. Finally, we compared beta_M-P_ between the MTV-based M and P subdivisions to evaluate the consistency between the stimulus selectivity of the BOLD responses and MTV-based parcellation of the LGN subdivisions.

### Test-retest reliability analysis

To assess the reproducibility of the MTV measurements and MTV-based LGN parcellation, we re-measured MTV of 13 subjects (mean age, 23.85 years; five females). The data acquisition and analysis of MTV retest data were identical to those of the main experiment. We evaluated the reproducibility of the MTV measurements within the LGN by calculating the inter-voxel correlation coefficient (R) between the test and retest data. In addition, we quantified the reproducibility of the MTV-based LGN parcellation by calculating the proportion of LGN voxels classified into the same subdivisions using the test and retest data. We evaluated the statistical significance of this proportion by comparing with a null distribution, which was obtained by shuffling the labeling of the M and P voxels 10,000 times and calculating the distribution of the proportions of voxels classified into the same subdivisions using the test and shuffled data. Finally, we also performed a comparison between the retest and BigBrain data using the same procedure as used for the test data.

## Results

We identified the whole LGN in individual subjects using PD weighted images (Viviano and Schneider, 2015) and then used MTV maps (Mezer et al., 2013) to identify the LGN subdivisions at the single-subject level. The validity of MTV-based parcellation of human LGN was evaluated by comparison with histological BigBrain data (Amunts et al., 2013) and with fMRI data collected from identical subjects. Furthermore, we tested the validity of LGN parcellation data obtained from other types of structural MRI images. Finally, we evaluated the test-retest reliability of the MTV-based parcellation of the human LGN.

### The LGN in PD weighted images

In each individual hemisphere, the position and shape of the whole LGN was visible in the PD weighted images (Figure 1A) as reported previously (Viviano and Schneider, 2015). We delineated the whole LGN in all individual hemispheres by manually inspecting the PD weighted images (Figure 1A, B; see Materials and Methods). Figure 1-S1 depicts the volume of whole LGN in all individual hemispheres, while the LGN volume identified from BigBrain histological data (Amunts et al., 2013) and previous structural MRI studies (Giraldo-Chica and Schneider, 2018; Mcketton et al., 2014) are also shown. Among the 15 subjects tested in this study, the mean (± s.e.m.) volume of the whole LGN was 153.48 mm^3^ ± 2.32 mm^3^ and 158.40 mm^3^ ± 1.83 mm^3^ in the left and right hemispheres, respectively. The LGN volume manually identified from PD weighted images was, on the whole, consistent with that obtained from BigBrain data (168.65 mm^3^ and 162.37 mm^3^ in the left and right hemispheres, respectively) and previous structural MRI images on the basis of similar PD weighted MRI data (Figure 1-S1; Giraldo-Chica and Schneider, 2018; Mcketton et al., 2014). The right LGN was marginally significantly larger than the left (*d’* = 0.608, *t_14_* = 2.136, *p* = 0.0508, paired *t*-test). Notably, the M and P subdivisions could not be identified through visual inspections of the PD weighted image.

### Identification of M and P subdivisions using the MTV fraction

Next, we analyzed the MTV maps (Mezer et al., 2013) co-registered with the PD weighted images for each individual subject (Figure 1C). In all of the individual hemispheres, we observed gradual changes in the MTV fractions within the LGN ROIs (Figure 1D); the superior-lateral part of the LGN exhibited a higher MTV fraction than the inferior-medial part. The distribution of MTV did not exhibit clear multiple peaks (Figure 1-S2), which made it difficult to determine a threshold value of MTV at which to separate the M and P subdivisions. The gradual MTV change seen within the LGN was likely due to the limited spatial resolution of the *in vivo* MTV measurements in this study (voxel size: 1 mm isotropic). We parcellated the human LGN by incorporating prior knowledge from an anatomical study demonstrating that the area of the P subdivision is roughly four times larger than that of the M subdivision (Andrews et al., 1997). Based on this knowledge, we classified the 20% of voxels with the lowest MTV as belonging to the putative M subdivision and the remaining 80% of voxels as belonging to the putative P-subdivisions (Figure 2; see Figure 2-S1 and 2-S2 for results obtained from other representative subjects; see Figure 3-S2 for results obtained using a different M/P ratio definition). As expected from the gradual difference in MTV fractions, the voxels classified as part of the putative M subdivision appeared at the inferior-medial part, whereas those classified as part of the putative P subdivision appeared at the superior-lateral part. We also note that these M and P subdivisions were observed as two distinct clusters of voxels which were highly continuous across slices in most hemispheres (Figures 2-S1 and 2-S2).

### Validations of M and P subdivisions identified using MTV data

We evaluated the validity of the MTV-based LGN parcellation by comparing it with the available histological data for human LGN subdivisions (Figure 2-S3). To do so, we calculated the centers of the coordinates among all of the M and P subdivision voxels classified based on the MTV in all hemispheres (Denison et al., 2014) and compared them with those from the BigBrain data (Figure 3; Figure 3-S1). In the BigBrain data, the center of the M subdivision was located in the medial, posterior, and ventral part of the LGN, while the center of the P division was located in the lateral, anterior, and dorsal part (dashed lines and open circles in Figure 3 and Figure 3-S1). We found that the center of the coordinates for the M and P subdivisions defined by the *in vivo* MTV data showed a similar trend as that seen using the BigBrain data, and this trend was well replicated across all subjects (solid lines and filled circles in Figure 3 and Figure 3-S1). Therefore, these results suggest that the LGN subdivisions identified using the MTV were in good agreement with the anatomical architecture defined using histological human LGN data.

**Figure 3:**
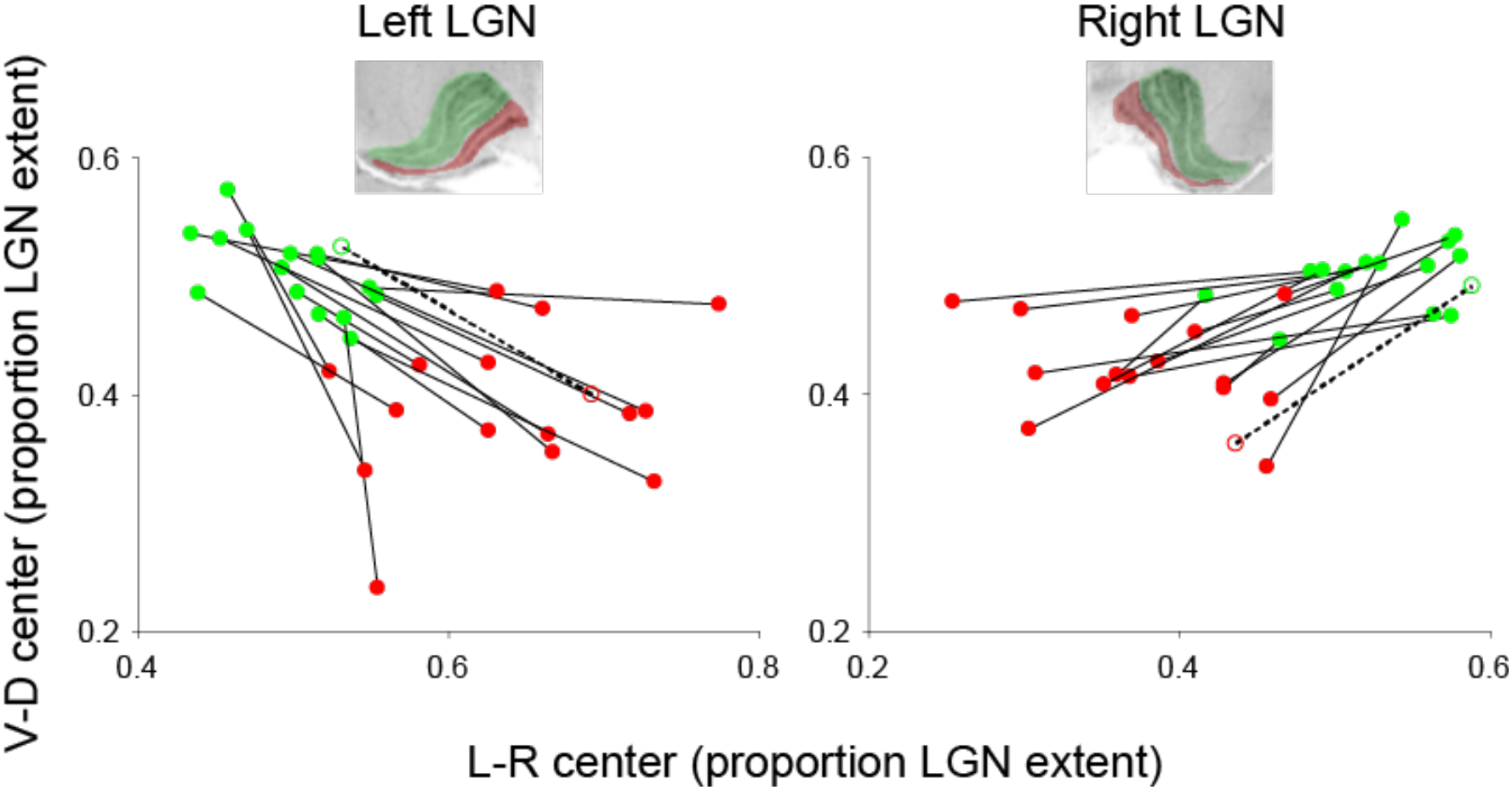
Center positions of the MTV-based M and P subdivisions compared with BigBrain data. The horizontal and vertical axes depict the center positions for the M (red) and P subdivisions (green; *left panel*, left LGN; *right panel*, right LGN). The horizontal and vertical axes represent the left-right and ventral-dorsal axes, respectively (Denison et al., 2014). The M and P subdivisions identified in a representative coronal slice using BigBrain data are inserted in each panel. The center positions were calculated in MNI coordinates and are plotted as a proportion of each subject’s entire LGN along a given axis. The filled circles and solid lines represent the spatial centers of the M and P subdivisions estimated from the MTV map. The open circles and dotted lines represent the spatial centers of the M and P subdivisions in a human histological dataset (BigBrain; Amunts et al., 2013). We found that the centers of the M voxels defined using the MTV were located more medially and ventrally than the P voxels in all hemispheres, which was consistent with the BigBrain histological data.

We replicated the MTV-based parcellation results using two different definitions of the ratio between the M and P subdivisions (i.e., M:P ratio = 19.0:81.0 or 28.9:71.1) to test whether the arbitrary choice of a fixed volume ratio used in the main analysis (1:4; Denison et al., 2014) affected the validity of the MTV-based parcellation. These two ratios correspond to the minimum and maximum proportions of M subdivisions across brains reported in a post-mortem study (Andrews et al., 1997) and determined by our investigations on the BigBrain data. We found that the overall pattern of MTV-based parcellation was well preserved in the current analysis, suggesting that when using a volume ratio from within the range reported by anatomical studies, MTV-based LGN parcellation is consistent with that found using histological data (Figure 3-S2).

### MTV-based M and P subdivisions exhibited different visual stimulus selectivities on fMRI

Using fMRI, we further tested whether the M and P subdivisions identified using the MTV exhibited the different visual stimulus sensitivities reported in previous macaque and human studies (Denison et al., 2014; Derrington and Lennie, 1984; Usrey and Reid, 2000; Zhang et al., 2016, 2015). We measured the BOLD responses to a pair of visual stimuli designed to activate the M and P subdivisions differently (Denison et al., 2014; Figure 4A; see *Materials and Methods, Functional MRI data acquisition*). We examined the extent to which the M and P subdivisions defined using the MTV exhibited different stimulus selectivity in their BOLD response.

**Figure 4:**
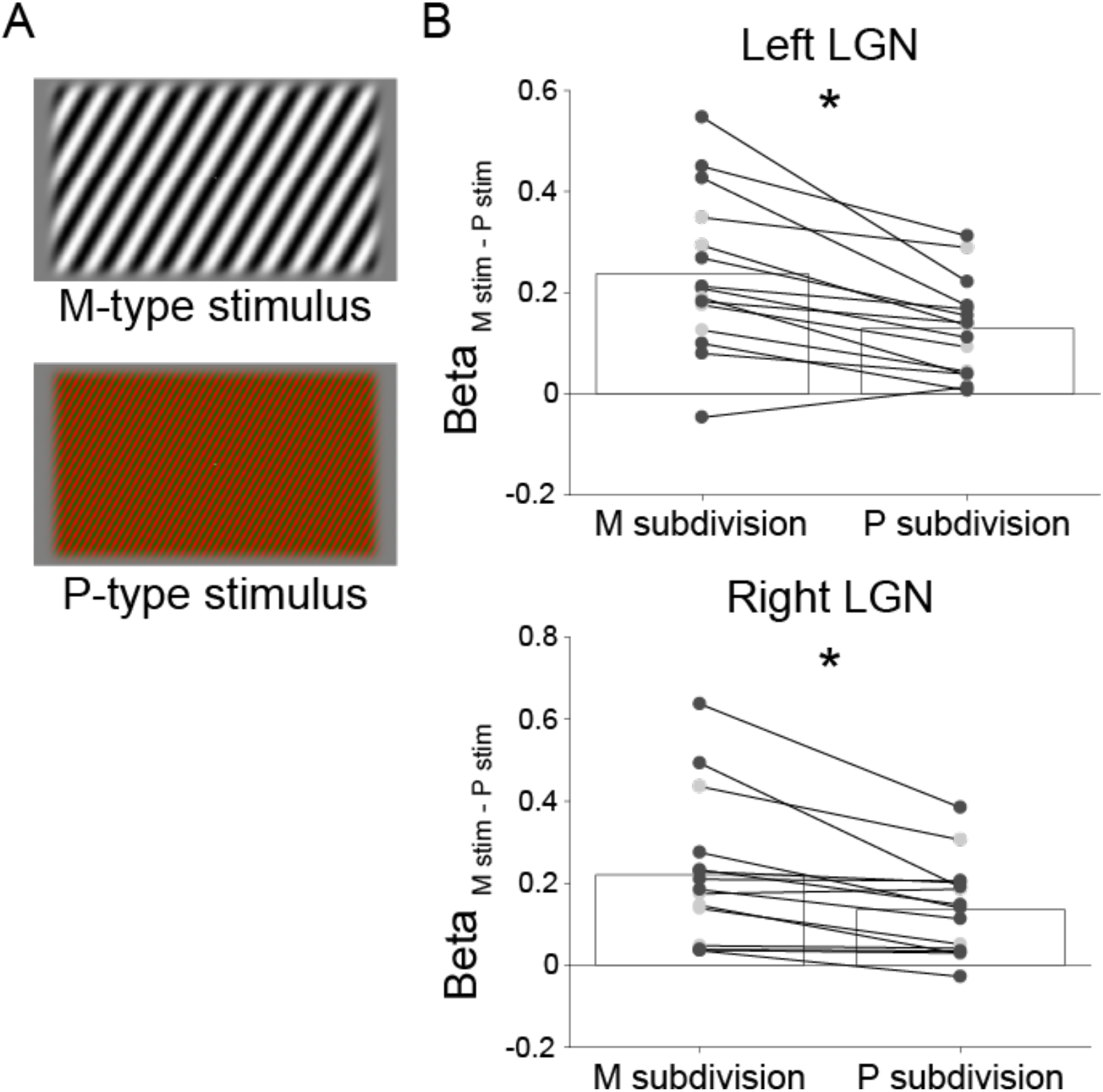
MTV-based M and P subdivisions exhibit stimulus selectivity as reported in previous literature. (A) Stimuli used to elicit differential BOLD responses from voxels with greater M subdivision representation and those with greater P subdivision representation. *Upper panel:* An achromatic, low spatial, and high temporal frequency with high luminance contrast grating stimulus used to activate the M subdivision. *Lower panel:* A high color contrast, high spatial, and low temporal frequency with low luminance contrast grating stimulus used to activate the P subdivision. These stimuli were adapted from Denison et al., (2014). See *Materials and Methods* for the details of the stimuli. (B) Stimulus selectivity measured by fMRI in the MTV-based M and P subdivisions (*top panel*, left LGN; *bottom panel*, right LGN; N = 15 for each). The vertical axis depicts the difference in the beta value between the M- and P-type stimuli (a positive value indicates more selective BOLD responses for the M-type stimuli). The dots indicate data in individual hemispheres. The dark and light gray dots represent the measurements with 1.5 mm isotropic (S1-S10) and 2.0 mm isotropic (S11-S15) voxels, respectively. The asterisks represent statistically significant differences in stimulus selectivity between the M and P subdivisions measured using the BOLD response (paired *t*-test, *: *p* < 0.005). Details of the fMRI methods are described in *Materials and Methods, Functional MRI data acquisition*.

We calculated the difference in the beta weights of the M- and P-type stimuli (Beta_Mstim-PStim_; a positive value indicated that the BOLD response was greater for the M-type stimuli) for the M and P subdivisions defined using the MTV in individual hemispheres (Figure 4B). A group analysis showed a significant difference in the Beta_Mstim-PStim_ between the M and P subdivisions (t_14_ = 4.5416, *p* = 0.0005 for the left hemisphere; t_14_ = 3.5838, *p* = 0.0030 for the right hemisphere, paired *t*-test). This consistency with known stimulus selectivity in the M and P subdivisions further supports the finding that MTV-based parcellation provides reasonable *in vivo* identification of LGN subdivisions at the level of individual hemispheres.

### Inter-subject variability and hemispheric differences in MTV fractions

We examined whether the MTV fractions in the estimated M and P subdivisions were consistent across the healthy subjects who participated in this study (Figure 5). The MTV fractions of the estimated M subdivision were 0.2525 ± 0.0032 and 0.2388 ± 0.0035 for the left and right hemispheres (mean ± S.E.M), respectively, whereas the MTV fractions of the estimated P subdivisions were 0.2923 ± 0.0029 and 0.2782 ± 0.0031 for the left and right hemispheres, respectively. Therefore, the inter-subject variability of the MTV fractions of each subdivision was much smaller than the mean difference between the M and P subdivisions in healthy subjects. Given the low variability in the measurements across the healthy population, MTV measurements of the LGN are reliable for use in evaluating how the LGN tissue in patients with, for example, eye disease deviates from that of control subjects.

**Figure 5:**
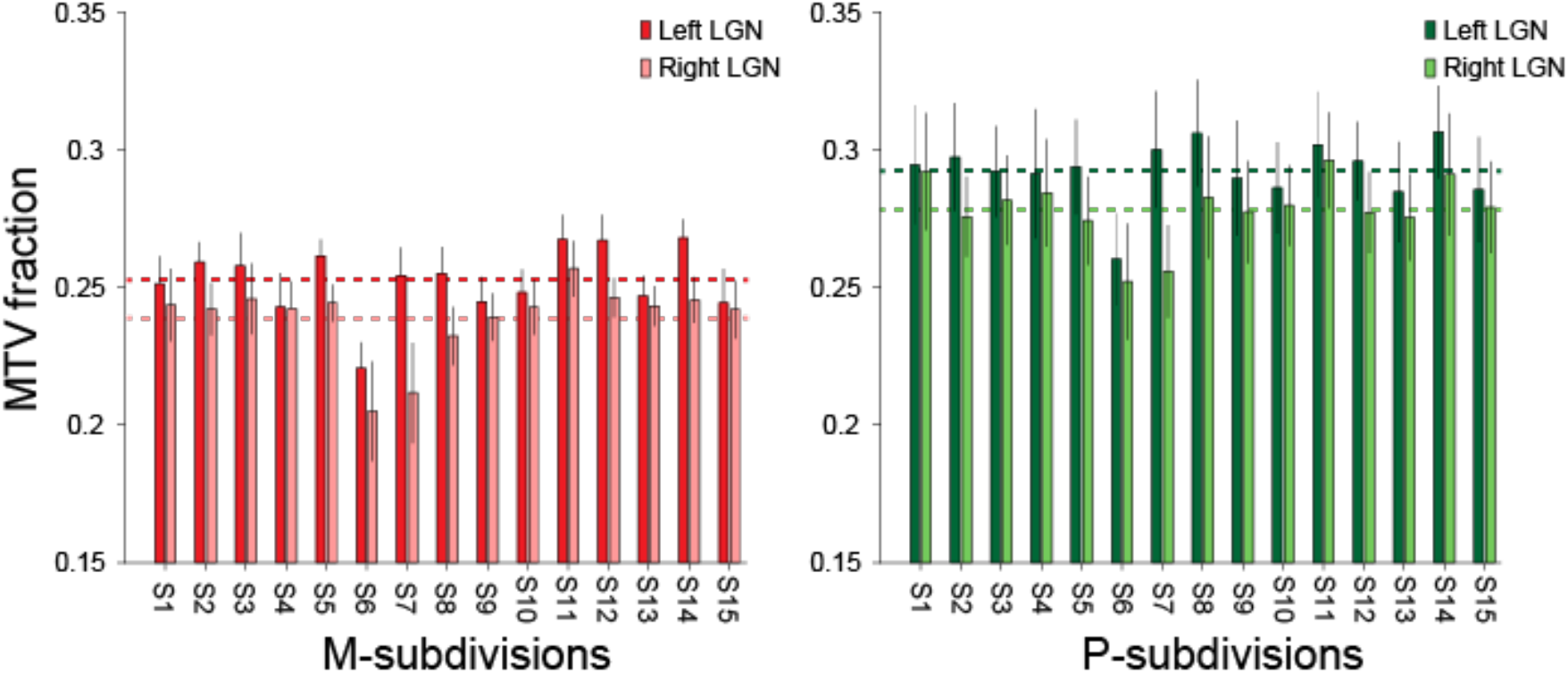
The MTV fractions in the estimated M and P subdivisions were consistent across all healthy subjects. The vertical axis depicts the MTV fraction averaged across voxels within the M (*left panel*, red) and P subdivisions (*right panel*, green) in individual hemispheres. The dark and light bars indicate the MTV fractions of the left and right hemispheres, respectively. The thin dotted lines indicate the averages for each hemisphere across subjects. The MTV fractions in these subdivisions were consistent across 15 healthy subjects. The error bars depict ±1 standard deviation across the voxels.

The MTV of the left LGN was found to be significantly higher than that of the right LGN in both the M and P subdivisions (mean MTV ± S.E.M for the M subdivision: 0.2525 ± 0.0032 and 0.2388 ± 0.0035 in the left and right hemispheres, respectively, *d’* = 1.0708, *t_14_* = 4.9078, *p* = 0.0002, paired *t*-test; P subdivision: 0.2923 ± 0.0029 and 0.2782 ± 0.0031 in the left and right hemispheres, respectively, *d’* = 1.220, *t_14_* = 5.1707, *p* = 0.0001, paired *t*-test). These significant hemispheric asymmetries in the MTV may not be due to measurement biases in the entire image, as no inter-hemispheric MTV differences were observed in the whole gray matter or white matter (mean MTV ± S.E.M: whole gray matter, 0.2038 ± 0.0015 and 0.2033 ± 0.0013 in the left and right hemispheres, respectively, d’ = 0.0933, *t_14_* = 0.6404, *p* = 0.5322; whole white matter, 0.3069 ± 0.0021 and 0.3073 ± 0.0022 in the left and right hemispheres, respectively, d’ = 0.0533, *t_14_* = 1.3652, *p* = 0.1937). The asymmetry of the MTV suggested that there may be some inter-hemispheric tissue differences in the LGN, although the exact microstructural interpretation of this effect requires further study.

### LGN M and P subdivisions were not parcellated from PD weighted images and T1w/T2w ratio maps

MTV is a useful method for obtaining quantitative measurements of brain tissue properties (Berman et al., 2018; Duval et al., 2017; Mezer et al., 2013). However, a number of studies have used other types of MRI-based metrics, such as the ratio between T1 weighted and T2 weighted images (T1w/T2w ratio), to evaluate tissue properties. The acquisition time for these images is shorter, but the measurements are not fully quantitative (Glasser et al., 2016; Glasser and Van Essen, 2011). Therefore, we tested whether the M and P subdivisions can be similarly parcellated using image intensities on non-quantitative structural MRI scans to evaluate the potential advantages of the MTV-based approach.

First, we tested LGN parcellation using the intensities of PD weighted images; these were used to identify the whole LGN (Figure 6A). Parcellation was also investigated using the T1w/T2w ratio map, which has been used often in previous studies to parcellate cortical areas (Glasser and Van Essen, 2011; Figure 6B). In both cases, we found that coordinates for the centers of the M and P subdivisions in the majority of subjects were inconsistent with those obtained using the histological data (BigBrain), suggesting that these maps were not sufficient for parcellating the M and P subdivisions in the LGN. This most likely occurred because these values are prone to measurement biases (Shams et al., 2019). The unsuccessful parcellation of the M and P subdivisions using the PD weighted images or T1w/T2w maps indicated the advantage of acquiring MTV data for LGN parcellation.

**Figure 6:**
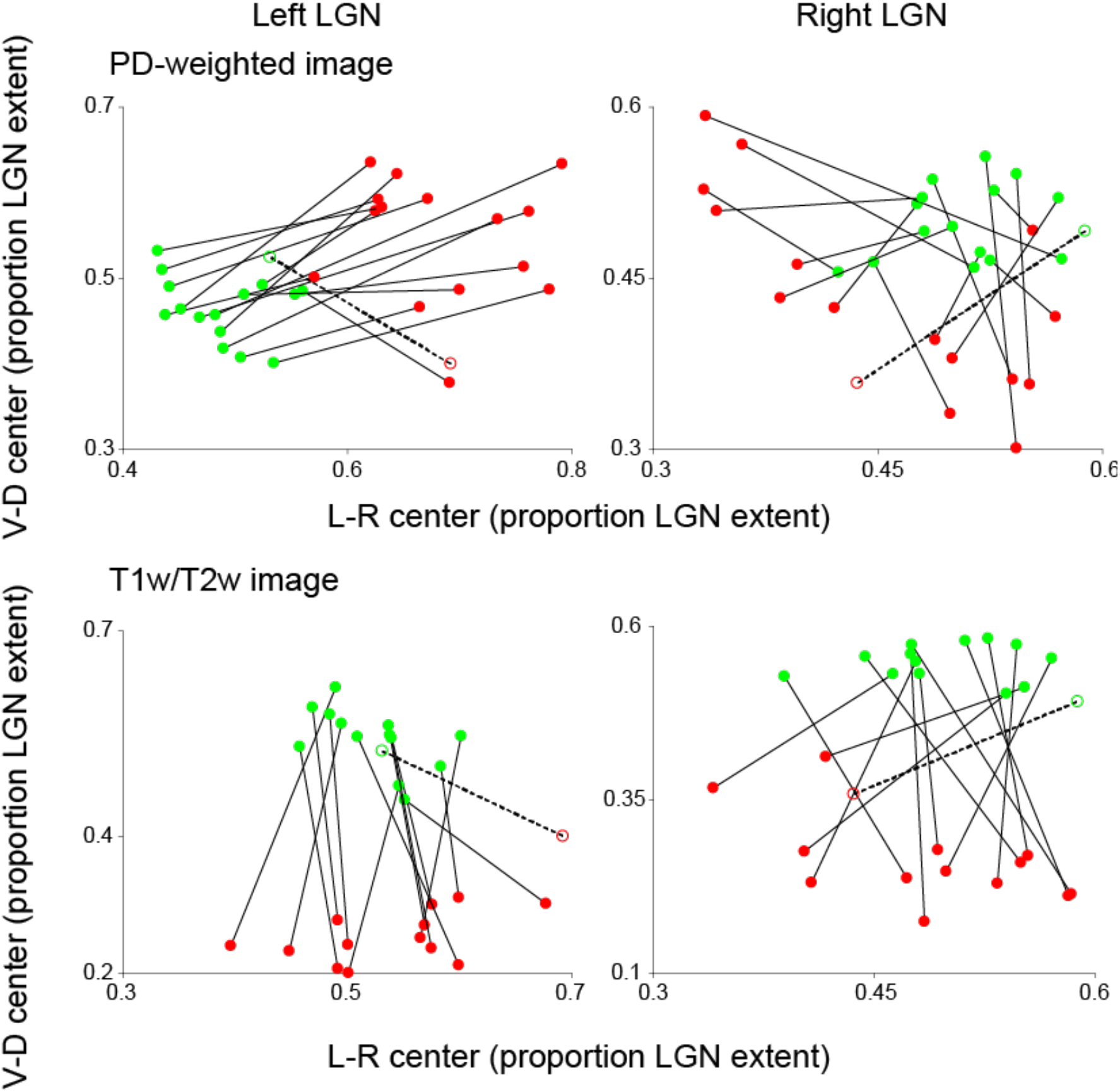
LGN subdivisions may not be identifiable from non-quantitative structural MRI maps. (A) LGN parcellation based on the image intensities of PD weighted images. The centers of the M and P voxels vary across the hemispheres and are inconsistent with the LGN coordinates obtained using the BigBrain data. (B) LGN parcellation based on the image intensities of T1w/T2w maps. In most hemispheres, the center points along the left-right axis were similar between the M and P subdivisions, unlike those seen using the BigBrain data. The conventions are identical to those used in Figure 3.

### LGN M and P subdivisions were parcellated using the MTV in the retest session

We tested the test-retest reliability of the M and P parcellations by performing the same MTV measurement in 13 subjects on a different day. The MTV fractions of voxels within the LGN ROI were highly correlated between the test and retest experiment (r = 0.78, Figure 7A). In calculating the probability that individual voxels can be classified into the same subdivisions between the test and retest experiments, we demonstrated that 85.62% and 82.31% of voxels in the left and right LGN, respectively, were classified in same subdivision in both the test and retest experiments (Figure 7B; mean across subjects). To assess the statistical significance of these numbers, we randomly classified 80% of voxels into the P subdivision and the remaining 20% into the M subdivision to obtain a null distribution. We repeated this process by shuffling the voxels 10,000 times. The maximum probabilities of the voxels being classified into the same subdivisions between the test and shuffled data were 71.73% and 71.55% for left and right hemispheres (mean across subjects), respectively, suggesting that the test-retest reliability of MTV-based parcellation was highly significant (*p* < 0.0001). Finally, using the retest dataset, we replicated the results indicating that the centers of the M and P subdivisions showed the same spatial patterns as in the histological data (Figure 7C). Taken together, these results support a considerable degree of reproducibility for our results from the MTV-based LGN parcellation.

**Figure 7:**
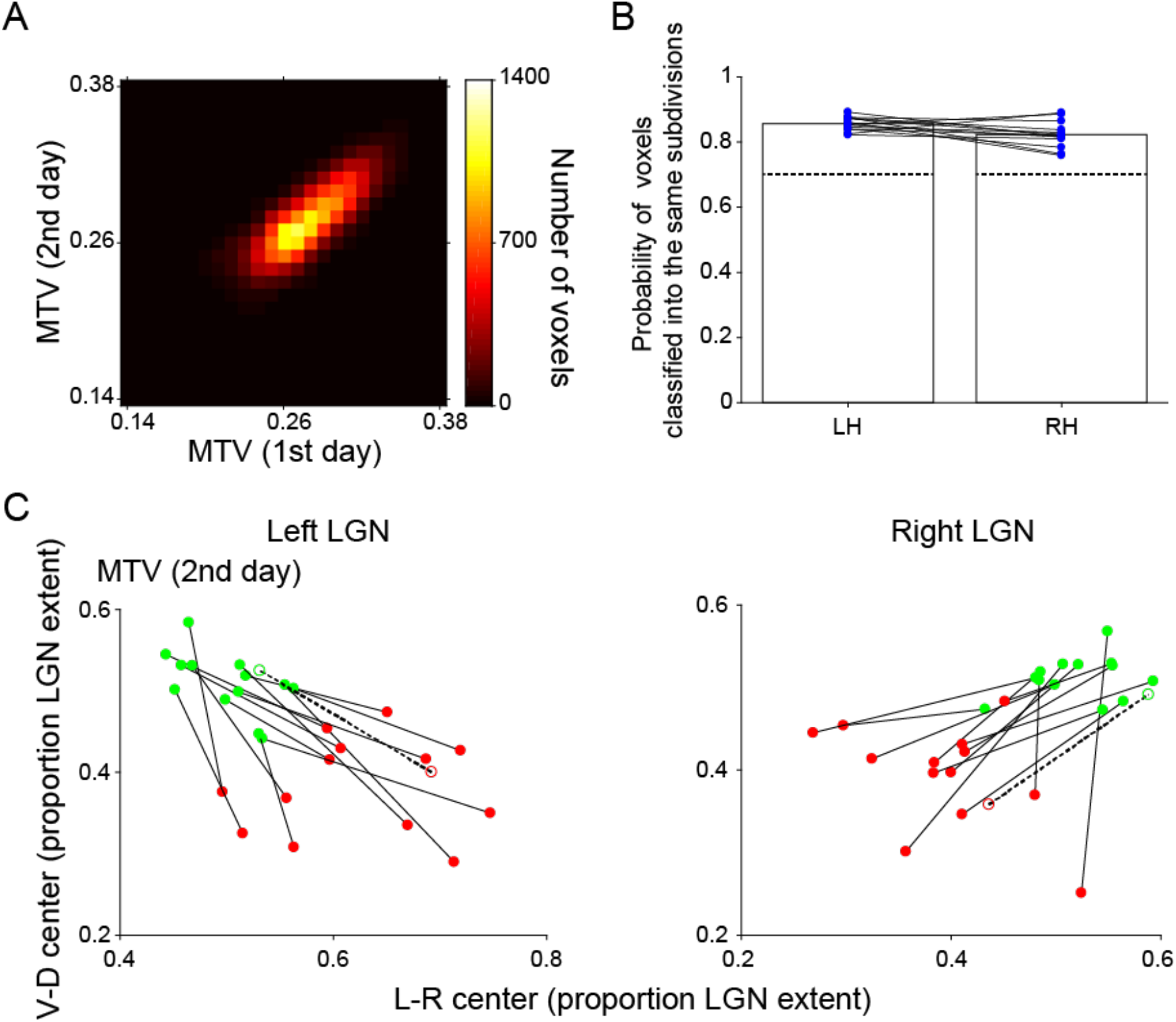
Test-retest reproducibility of MTV-based parcellation. (A) Two-dimensional histogram comparing the MTV measurements across days in LGN voxels. The data are derived from LGN voxels pooled across subjects, who took part in the retest scans (N = 13). The color map indicates the number of voxels. The correlation coefficient of MTV measurements across days was 0.78 (*p* < 10^−10^). This high correlation coefficient indicated that the MTV measurements were reproducible across days. (B) Reproducibility of classification. The vertical axis depicts the probability that individual voxels were classified into the same LGN subdivisions between the test and retest dataset. The individual dots depict the results in individual hemispheres. The dotted lines depict the maximum probability of voxels being classified into the same subdivisions between the test and shuffled data, which was equal to *p* = 0.0001. (C) The centers of the coordinates for the M and P subdivisions identified with MTV-based parcellation using retest dataset (filled circles and solid lines). We replicated the results on the centers of M and P subdivisions in a consistent manner with those in BigBrain data (Amunts et al., 2013; open circles and dotted line). The conventions are identical to those used in Figure 3.

## Discussion

It is widely known that the human LGN consists of functionally and anatomically different subdivisions. However, identifying these subdivisions in individual living human brains using conventional structural neuroimaging methods had been challenging. In this study, we demonstrated the approximate parcellation of LGN subdivisions at the single-subject level by combining *in vivo* structural MRI methods (multiple PD weighted imaging and MTV measurement). The spatial positions of the identified LGN subdivisions were consistent with those identified using histological data (Amunts et al., 2013). Furthermore, using fMRI, we confirmed that these subdivisions exhibit different stimulus selectivity, which is consistent with the findings of previous physiological studies. Finally, we confirmed that MTV-based LGN parcellation is highly consistent across datasets acquired on different days, suggesting that the proposed method is highly reproducible. Other non-quantitative MRI methods did not provide LGN parcellation consistent with histological data. Taken together, this study provides evidence of the utility of this quantitative structural MRI approach to LGN parcellation and establishes methods of measuring the structural properties of human LGN subdivisions in single living human subjects using a clinically feasible 3T MRI scanner.

### Microstructural origin of MTV-based parcellation

Our results demonstrated that the MTV fraction can be a useful measurement for distinguishing M and P subdivisions. One might ask what types of microstructural differences lead to MTV differences between M and P subdivisions. In principle, the MTV quantifies non-water macromolecular volumes on the basis of calibrated quantitative PD maps (Mezer et al., 2013). Phantom experiments confirmed that MTV measurements correlate with the lipid fraction (Filo et al., 2019; Mezer et al., 2013; Shtangel and Mezer, 2020). However, there is no established theory on how much MTV variance in a particular brain area can be explained by specific types of microstructural properties. A number of histological studies on non-human primates reported differences in anatomical properties between M and P subdivisions, such as the higher neuronal cell density (Hassler, 1966; Yücel et al., 2003, 2000) and greater myelin content in the P subdivision (Pistorio et al., 2006). A recent post-mortem human study confirmed these properties (Müller-Axt et al., 2020). These results are in line with our results showing a larger MTV fraction in the P subdivision than in the M subdivision, since both a larger number of cells and greater myelin content will result in a larger lipid volume fraction. It is also possible that other neurobiological factors such as glial cell density also partly explain the difference in MTV between the M and P subdivisions. This remains an open question for future investigations that more directly compare histology and quantitative MRI maps.

In this study, we also determined a significant difference in the MTV fraction between the left and right LGN. To our knowledge, this is the first neuroimaging study to demonstrate this inter-hemispheric difference in human LGN tissue properties. While speculative, these results may be related to those of a previous study showing a difference in the number of neurons in left and right LGN of rhesus monkeys (Williams and Rakic, 1988). Further histological examination will be necessary to establish what type of microstructural differences exist between the hemispheres of the human LGN.

### Advantage of MTV-based parcellation over other structural MRI methods

In standard practice, many neuroimaging studies have utilized T1-weighted and/or T2-weighted images to locate cortical areas or subcortical nuclei. While the relative values in these images are useful for identifying the borders between gray and white matter, their absolute values cannot be interpreted as being quantitative, because the measurements are affected by multiple sources of inhomogeneity such as B1 transmitter (B_1_^+^) inhomogeneity or coil gain bias. Recent developments in quantitative MRI have enabled the quantification of MRI parameters, which allows the comparison of brain tissue properties between human subjects (Cercignani et al., 2018; Forstmann et al., 2016; Keuken et al., 2017; Mezer et al., 2013; Weiskopf et al., 2015). These quantitative MRI measurements have provided valuable insights into the tissue properties of cortical areas (Lutti et al., 2014; Sereno et al., 2013) and white matter (Stüber et al., 2014; Takemura et al., 2019).

Mezer and colleagues (2013) proposed MTV methods and demonstrated consistency of MTV measurements with lipid volume fractions in a phantom, high test-retest reproducibility, and sensitivity for white matter tissue changes in patients with multiple sclerosis. A strong advantage of this method is its independence from the static magnetic field strength, since it is based on PD measurements calibrated by assuming that the water fraction in CSF voxels is 100%. In fact, Mezer et al. (2013) demonstrated that MTV measurements in the brain are consistent across measurements performed using different types of hardware. Therefore, we chose MTV mapping as a potential method for parcellating the human LGN, since it is relatively independent from hardware choices and thus useful for future clinical studies.

We found that MTV enabled the parcellation of the LGN in a consistent manner to that seen using histological data (Figure 3) and subdivisions identified by MTV exhibited the stimulus selectivity consistent with previous physiological studies (Figure 4). MTV-based parcellation was superior to parcellation based on non-quantitative MRI maps (PD weighted images or T1w/T2w maps; Figure 6). This is most likely because the MTV was corrected for B_1_^+^ inhomogeneity, while the other maps were not. While the T1w/T2w map has been demonstrated to enhance tissue contrast and thus be useful for delineating borders between brain areas (Glasser and Van Essen, 2011) and has advantages in terms of shorter acquisition time, several studies demonstrated inconsistencies between T1w/T2w and quantitative MRI measurements that were more sensitive to myelin (Arshad et al., 2017; Hagiwara et al., 2018; Uddin et al., 2018). These inconsistencies are most likely due to the fact that T1w/T2w images are not calibrated for B_1_^+^ inhomogeneity (Glasser and Van Essen, 2011). While we have not excluded the possibility that ad-hoc B_1_^+^ bias field correction (Glasser et al., 2013) may improve LGN parcellation using T1w/T2w images, our results demonstrated that MTV-based LGN parcellation performed better than that with T1w/T2w, most likely because of the superior calibrations for B_1_^+^ inhomogeneity in the LGN.

### Comparison with fMRI-based LGN parcellation

A few fMRI studies have examined the spatial pattern of visually-evoked BOLD signals in the LGN (Denison et al., 2014; Zhang et al., 2015). These studies demonstrated that clusters of LGN voxels preferentially respond to distinct types of visual stimuli, which was consistent with neurophysiological findings, suggesting that an approximate identification of the LGN subdivisions in living humans can be achieved using fMRI-based measurements of visual stimulus sensitivities. Quantitative structural MRI-based parcellation methods as shown in this study have several advantages compared with fMRI-based parcellation methods. First, the fMRI-based methods require a precise control of visual stimuli, which involves the presentation of isoluminant stimuli (Denison et al., 2014); this is unnecessary when using the structural MRI-based methods. Furthermore, the use of visual stimuli limits the application of LGN parcellation methods in clinical studies of patients with visual field loss or when using MRI scanners without visual stimulus presentation equipment. Second, structural MRI-based methods are more spatially precise, since the voxel size for MTV measurements (e.g., 1 mm isotropic in this study) is generally smaller than those used in fMRI experiments (e.g., 1.8 × 1.8 × 1.5 mm for 3T or 1.2~1.5 mm isotropic for 7T in Denison et al., 2014). When using fMRI, large veins passing through multiple voxels can limit the spatial specificity of the BOLD signal (Kay et al., 2019; Uludağ and Blinder, 2018). Therefore, our methods are advantageous in terms of spatial precision, although fMRI-based methods are sure to improve (Huber et al., 2018, 2014; Kay et al., 2020). Finally, MTV-based parcellation has higher test-retest reliability across days (Figure 7) compared with that reported previously in an fMRI study (r < 0.4; Denison et al., 2014). Therefore, MTV-based parcellation provides more stable identification of M and P subdivisions in individual living human brains.

### Limitations and future directions

In this study, we classified voxels into M and P subdivisions using a fixed volumetric ratio (1:4). However, previous studies reported inter-subject variability in the ratio of the M and P subdivision volumes in post-mortem human data (Andrews et al., 1997; Hickey and Guillery, 1979). Therefore, it would be more ideal to classify the LGN voxels into M and P subdivisions by fitting a mixture model composed of two curves with distinct peaks to the distribution of the MTV fractions in each individual LGN. However, this approach was not practical in this study as the MTV-based distribution of LGN voxels in our *in vivo* data did not show two distinct peaks corresponding to the M and P subdivisions (Figure 1-S2). Given that a recent study using high-resolution *ex vivo* quantitative structural MRI (220 μm isotropic) using 7T MRI demonstrated a parcellation of LGN subdivisions using the aforementioned curve fitting procedure (Müller-Axt et al., 2020), we expect that future improvement in the spatial resolution of *in vivo* quantitative MRI will provide LGN parcellation without assuming a fixed volumetric ratio, enabling quantitative comparisons of M and P subdivisions volumes across subjects.

The other limitation of the MTV-based parcellation method proposed in this work was the relatively long acquisition time. Our method of identification required the acquisition of multiple structural images, each of which required a considerable length of time to obtain (60–90 and 28 min for the PD weighted image and quantitative structural MRI, respectively) to ensure higher signal quality at the LGN and quantification of the MTV (Giraldo-Chica et al., 2015; Giraldo-Chica and Schneider, 2018; Mcketton et al., 2014; Mezer et al., 2013; Viviano and Schneider, 2015). While the trade-off between signal quality and acquisition time is inherent to any neuroimaging method, prolonged acquisition times limit the application of MTV-based parcellation to studies on clinical populations. Further development of the quantitative MRI method with shorter acquisition times (Caan et al., 2019; Ma et al., 2013; Marques et al., 2010; Warntjes et al., 2008) is essential to increasing the opportunity to measure LGN subdivisions in a wide range of populations.

Thin koniocellular layers (K layers) are known to exist between each layer of the M and P subdivisions in the LGN (Guillery and Colonnier, 1970). Previous histological studies have indicated that the K layers have distinct anatomical properties compared with those in the M and P subdivisions (Hendry and Reid, 2000). Considering the location of K layers, the MTV fractions in many LGN voxels are likely to be affected by partial volumes between the K layers and M or P subdivisions. Therefore, part of the variance in the structural measurements may be affected by the anatomical properties of the K layers.

Despite the limitations discussed above, this study provides an important step toward establishing a method of parcellating human LGN subdivisions and quantifying their tissue properties in living human brains. Once the limitation of acquisition time has been overcome, MTV-based *in vivo* human LGN measurements will create fruitful opportunities to investigate how the properties of the human LGN are related to brain functions and affected as a consequence of disease. For example, psychophysical studies have suggested that certain properties of the M subdivision may be crucially related to reading performance (Chase and Jenner, 1993; Demb et al., 1998; Felmingham and Jakobson, 1995; Main et al., 2014; Stein, 2001). Also, psychophysical investigations of glaucoma patients have suggested that the M subdivision may be damaged earlier than P subdivision (Cello et al., 2000; Maddess et al., 1992). Extensions of this study will create a powerful tool for directly comparing LGN tissue properties with psychophysical performance to improve our understanding of how various properties of the LGN subdivisions are related to visual functions and disorders.

---

**Figure 1-S1:**
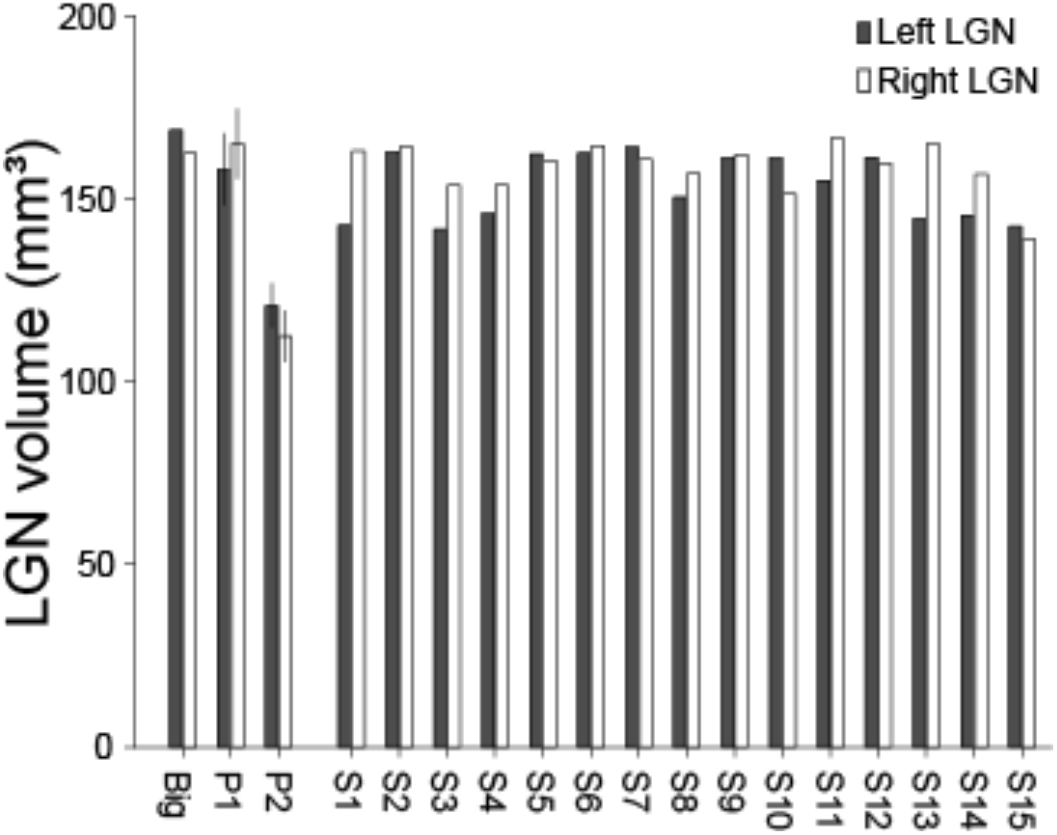
The LGN volume estimated in the current study and previous anatomical and neuroimaging studies. The gray and white bars depict the LGN volumes of the left and right hemispheres, respectively. The LGN volumes in this study (N = 15, from S1-S15) are within the range of the BigBrain (Big, left end) and previous structural MRI studies (P1, Mcketton et al., 2014 and P2, Giraldo-Chica et al., 2018). We note that both the P1 and P2 data were taken from data collected from healthy control subjects in each paper. Error bars depict ±1 S.E.M.

**Figure 1-S2:**
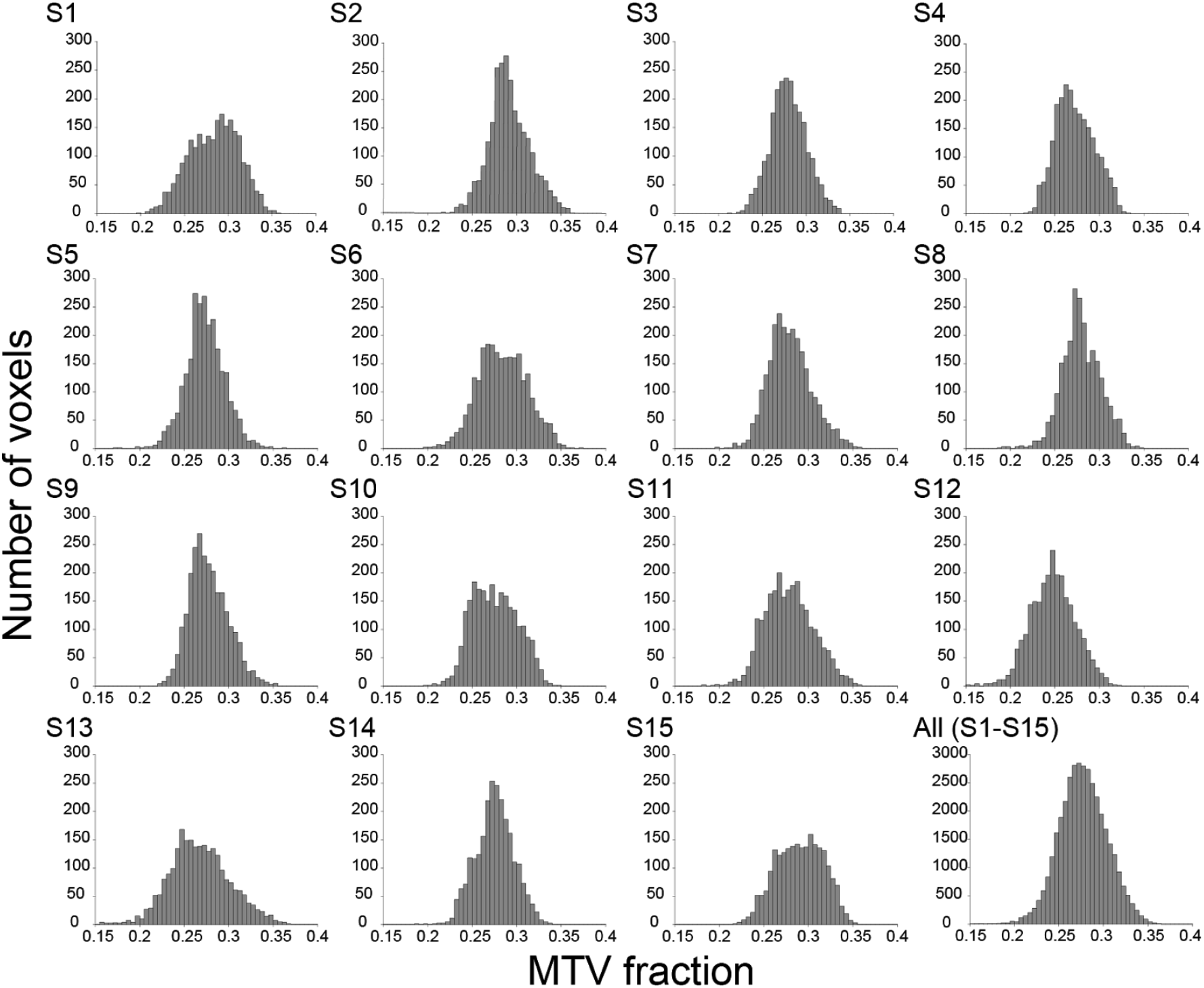
A histogram of the MTV fraction in all LGN voxels for all subjects (S1-S15) and those pooled across subjects (All). The horizontal axis represents the MTV fraction and the vertical axis represents the number of voxels. Bin widths are 0.005. We did not find a visible bimodal distribution of the MTV fraction within the LGN ROI.

**Figure 2-S1:**
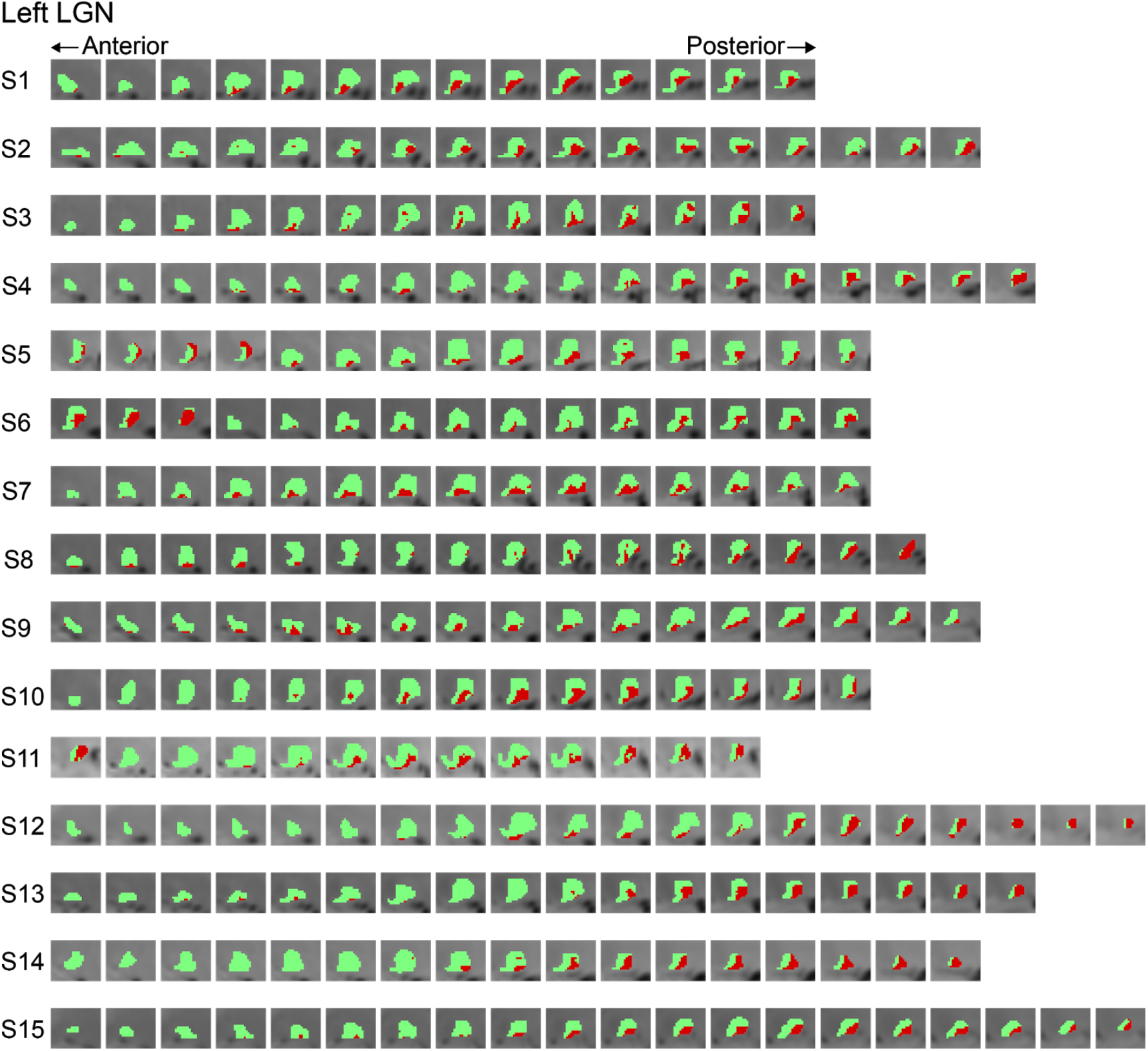
LGN subdivisions parcellated using the MTV in the left hemispheres of all subjects. The M and P subdivisions identified by the MTV in all subjects (N = 15) were overlaid on a series of coronal sections from PD weighted images (left: anterior section; right: posterior section; distance between sections: 0.5 mm). The conventions are identical to those used in Figure 2B.

**Figure 2-S2:**
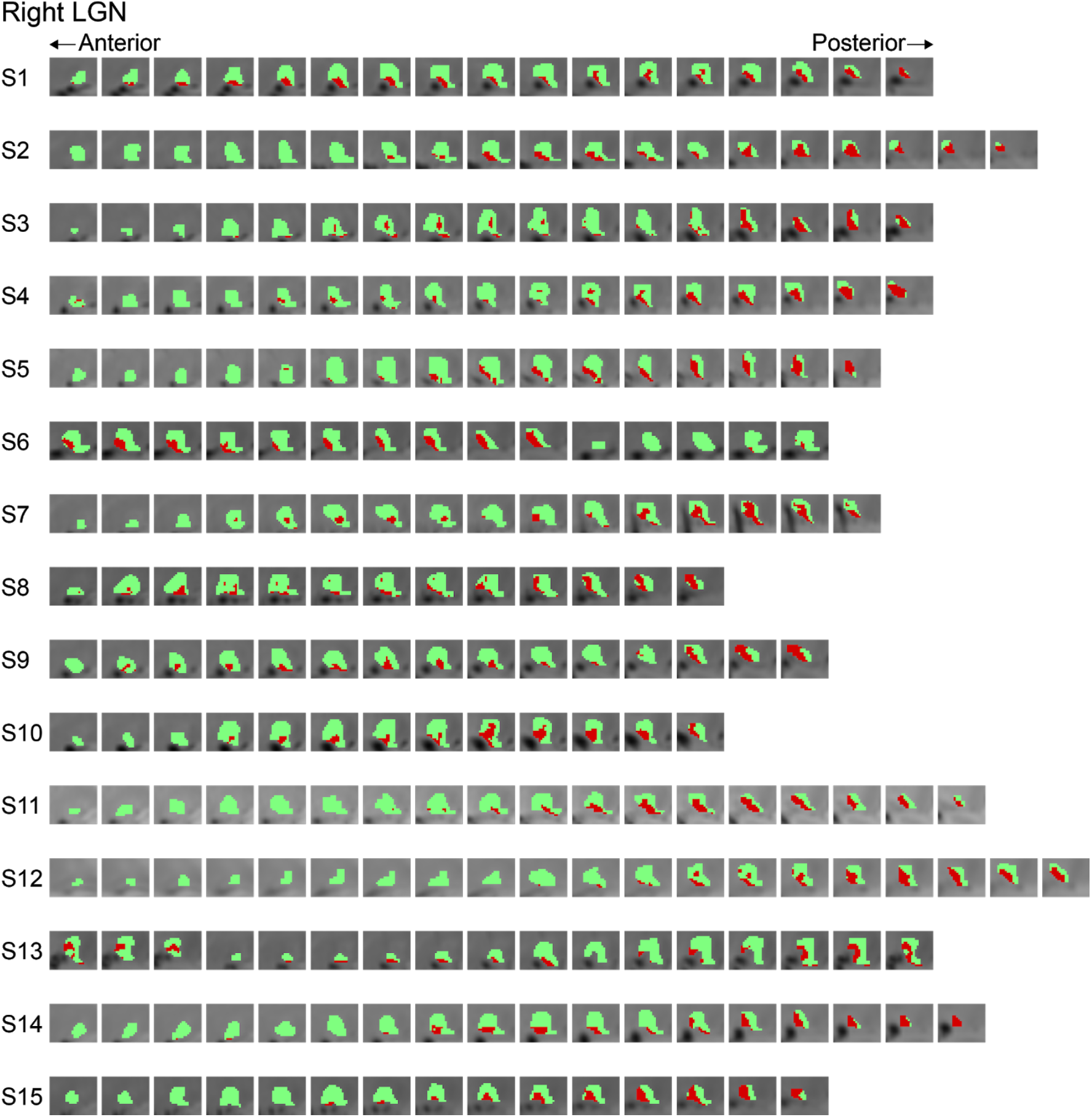
LGN subdivisions parcellated using the MTV in the right hemispheres of all subjects. The conventions are identical to those used in Figure 2B and 2-S1.

**Figure 2-S3:**
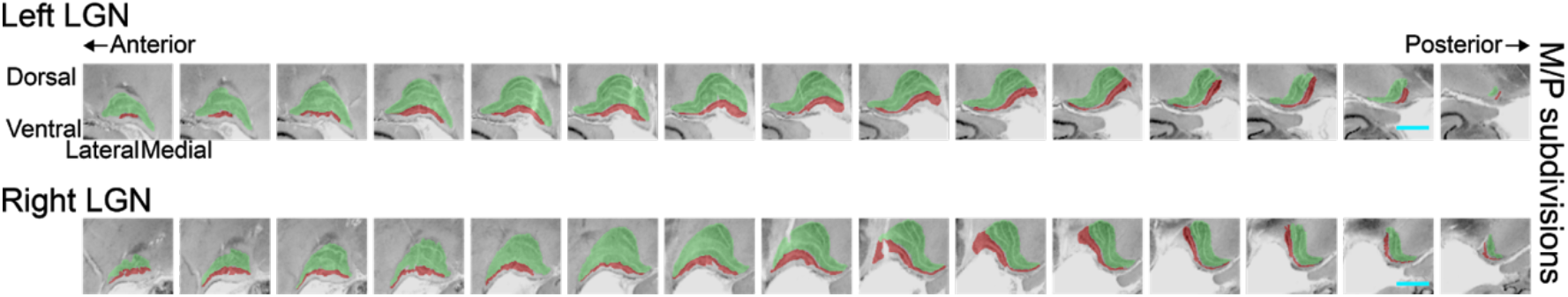
LGN subdivisions on a series of coronal sections from BigBrain data. M (translucent red) and P subdivisions (translucent green) manually identified from BigBrain data (upper panel, left hemisphere; lower panel, right hemisphere). The cyan scale bar indicates 4 mm, and the distance between the sections was 0.5 mm. All other conventions are identical to those used in Figure 2.

**Figure 3-S1:**
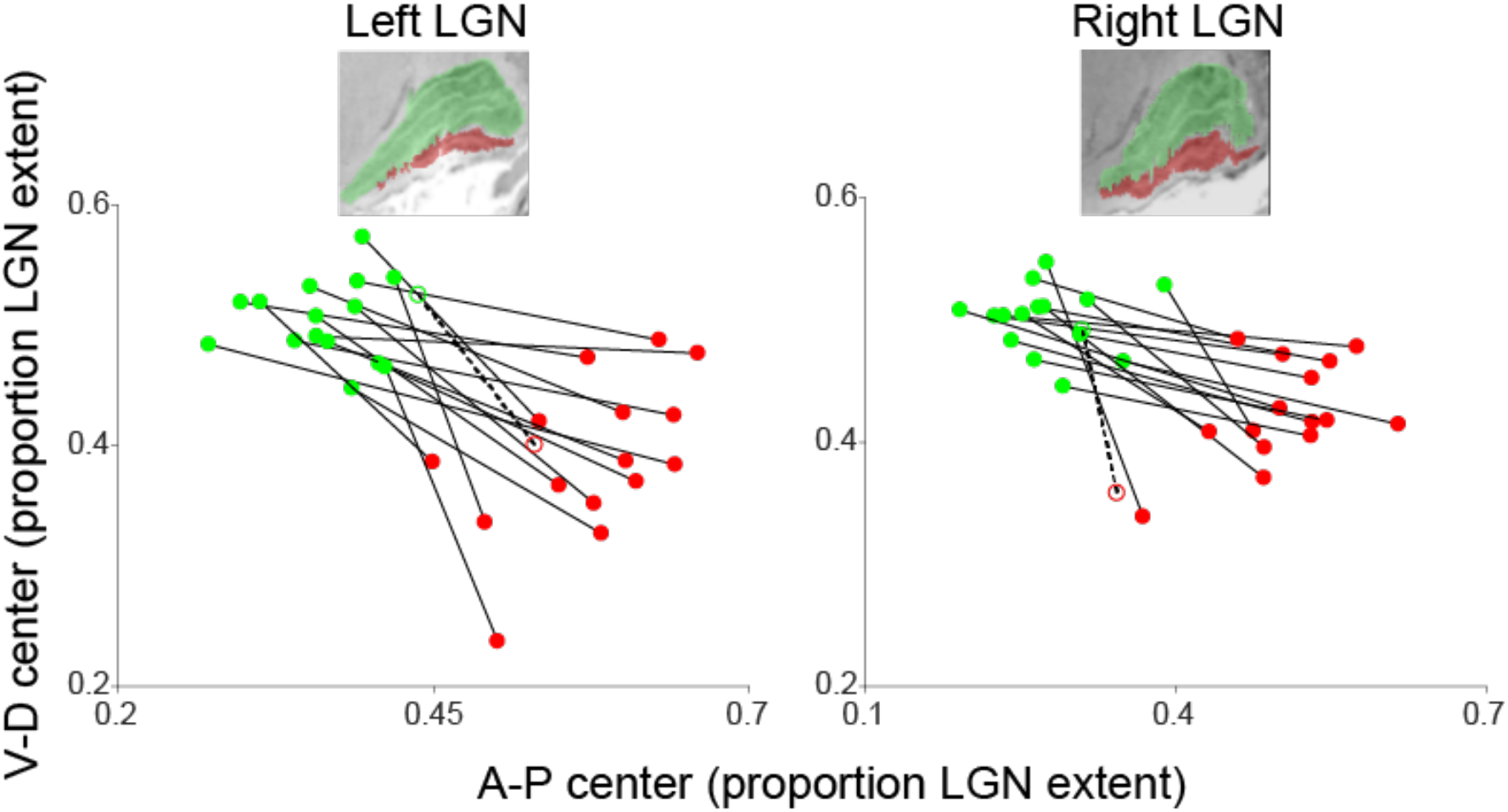
Center positions of the MTV-based M and P subdivisions compared with BigBrain data in the anterior-posterior/ventral-dorsal axis. The horizontal and vertical axes represent the anterior-posterior and ventral-dorsal axes, respectively. The M and P subdivisions identified in a representative sagittal slice using BigBrain data are inserted in each panel. All other conventions are identical to those used in Figure 3.

**Figure 3-S2:**
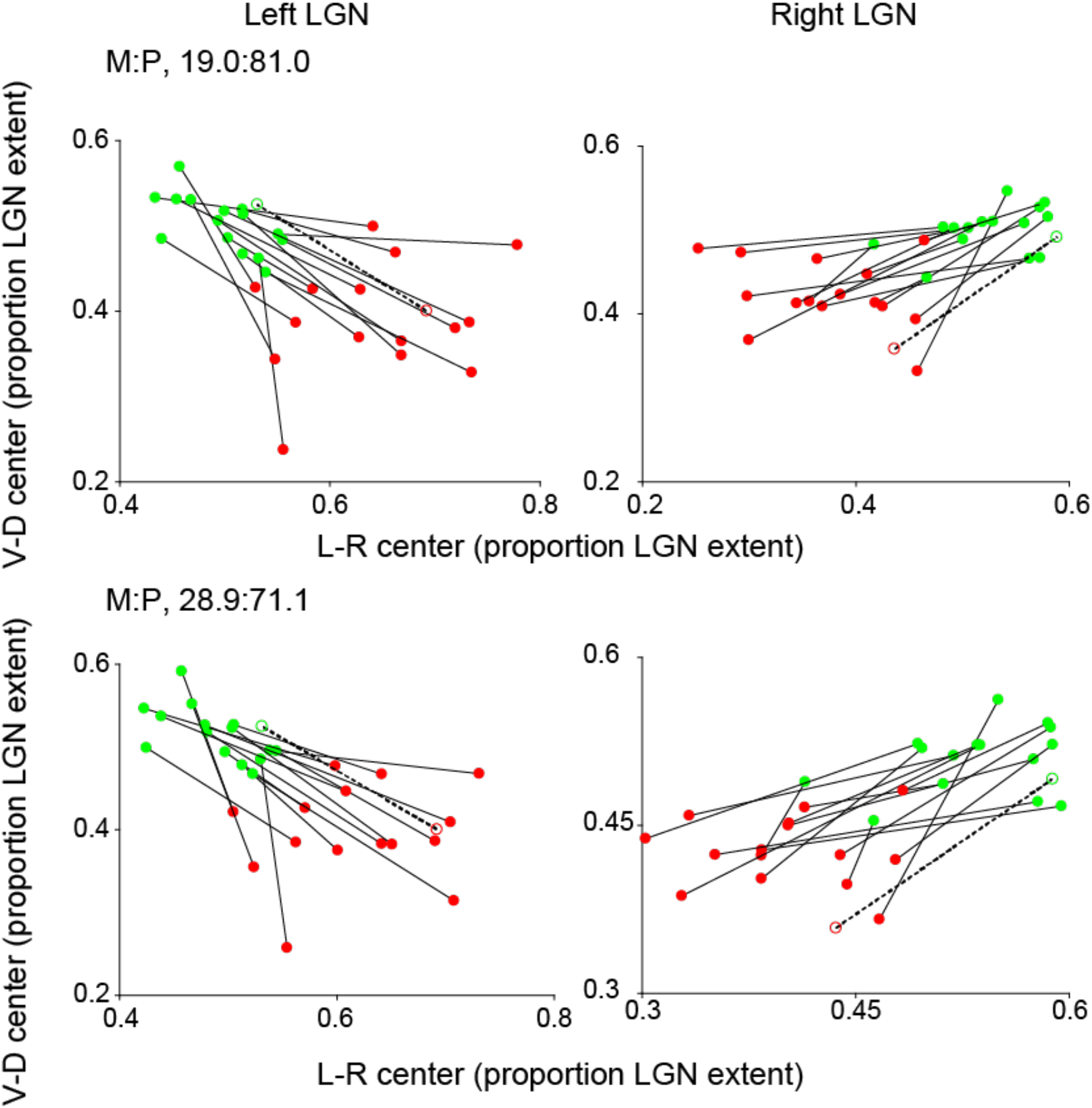
MTV-based parcellation was robust against the ratios of M and P subdivisions. The plots depict the centers of the M and P subdivisions estimated by the MTV fractions with different M:P ratios from the main analysis (top panels, M: 19.0%; bottom panels, M: 28.9%). These two ratios correspond to the minimum and maximum proportions of the M subdivision across brains reported in a previous post-mortem study (Andrews et al., 1997) and our investigations on the BigBrain data (Amunts et al., 2013). The conventions are identical to those used in Figure 3.

## Acknowledgments

We thank Yusuke Sakai for supporting the acquisition and analysis of the data.

